# Switching ON Hydrogen Sulfide: A Chemogenetic Toolkit for Spatially Resolved H_2_S Manipulation

**DOI:** 10.1101/2025.05.12.653628

**Authors:** Asal Ghaffari Zaki, Seyed Mohammad Miri, Emre Vatandaşlar, Sven Vilain, Esra Nur Yiğit, Mehmet Şerif Aydın, Muhammed İkbal Alp, Emrah Eroglu

## Abstract

Hydrogen sulfide (H_2_S) is emerging as a multifaceted signalling molecule that shapes energy metabolism, vascular tone, cancer biology and neurodegeneration. Although fluorescent and genetically encoded sensors now allow real-time H_2_S imaging, tools for selective control of intracellular H_2_S remain scarce. Here we benchmark two substrate-based chemogenetic enzymes—the yeast D-amino-acid oxidase (mDAAO) and the *Salmonella typhimurium* D-cysteine desulfhydrase (stDCyD)—for controlled H_2_S production. Both enzymes catalyse the conversion of D-cysteine to H_2_S, yet only mDAAO concomitantly generates hydrogen peroxide, introducing an unwanted oxidative signal. In contrast, stDCyD produces exclusively H_2_S, thereby offering a clean and efficient means to elevate intracellular H_2_S *in vitro*. Our data position stDCyD as the superior chemogenetic actuator for dissecting H_2_S biology without confounding redox artefacts.

## Introduction

Hydrogen sulfide (H_2_S) has emerged as a critical gasotransmitter alongside nitric oxide (NO) and carbon monoxide (CO), with growing recognition of its diverse roles in physiological and pathological processes^1,2^. Originally regarded as merely a toxic gas, H_2_S is now known to be endogenously produced in mammalian cells by enzymes such as cystathionine β-synthase (CBS), cystathionine γ-lyase (CSE), and 3-mercaptopyruvate sulfurtransferase (MPST)^3^, regulating key biological functions ranging from vascular tone^4^ to mitochondrial bioenergetics^3,5^ and neuromodulation^6^. Dysregulation of H_2_S homeostasis has been implicated in a range of disorders, including cardiovascular disease^7^, cancer^8^, and neurodegenerative conditions such as Alzheimer’s disease^9^.

Despite these advances, our ability to study H_2_S dynamics at the cellular level remains significantly limited. Traditional approaches to modulate intracellular H_2_S levels largely rely on genetic manipulations—such as the overexpression or knockout of H_2_S-producing enzymes^10^— or pharmacological interventions, including the administration of sulfide-releasing salts like sodium hydrosulfide (NaHS) or slow-releasing H_2_S donors^11^. However, these methods suffer from major drawbacks, including poor spatiotemporal resolution, lack of reversibility, and off-target effects that complicate the interpretation of results. Particularly in single-cell studies or in complex tissue environments, the inability to achieve precise, real-time control of H_2_S concentrations limits the scope of mechanistic investigations.

In recent years we have witnessed the development of chemical probes and genetically encoded sensors for real-time detection of H_2_S^12–15^. These advances have provided valuable insights into the spatial and temporal dynamics of H_2_S signaling. However, detection alone is insufficient for functional studies; tools are needed that not only report on H_2_S levels but also enable active manipulation with high precision.

A novel chemogenetic approach for intracellular H_2_S manipulation would overcome the limitations of exogenous donor administration, providing unprecedented spatiotemporal control and the possibility to dissect causality in H_2_S signaling pathways.

In this study, we introduce and characterize two enzymes with potential for use as chemogenetic tools to manipulate intracellular H_2_S levels. We investigated a modified D-amino acid oxidase (mDAAO) and Salmonella typhimurium derived D-cysteine desulfhydrase (stDCyD) that, in the presence of D-cysteine, can generate H_2_S^16,17^.

Beyond technical innovation, the ability to precisely manipulate H_2_S levels will have broad implications for understanding H_2_S signaling in both health and disease. In the vascular system, H_2_S is a known vasodilator modulating endothelial nitric oxide synthase (eNOS)^18,19^. In cancer, altered H_2_S metabolism contributes to tumor progression, angiogenesis, and chemoresistance^20^. In the nervous system, H_2_S acts as a neuromodulator influencing synaptic transmission and neuroprotection^6^, but its dysregulation is associated with neurodegenerative diseases^21,22^. Thus, the development of a chemogenetic system for H_2_S manipulation is poised to fill a critical gap in the field, enabling novel experimental paradigms that were previously inaccessible. By introducing a robust, targeted, and highly specific strategy to control intracellular H_2_S concentrations, we provide a powerful tool to unravel the complex biology of this important gaseous signaling molecule.

## Results

Figure 1 illustrates the distinct pathways by which the mDAAO and stDCyD enzymes mediate H_2_S production within cells. stDCyD, a pyridoxal phosphate (PLP)-dependent enzyme, catalyzes the direct conversion of D-cysteine into H_2_S, with pyruvate generated as a byproduct^1*7*^. In contrast, mDAAO first oxidizes D-cysteine to its corresponding α-keto acid, namely mercaptopyruvate. Subsequently, mercaptopyruvate can be further metabolized to H_2_S by the endogenous enzyme mercaptopyruvate sulfurtransferase (MPST)^16,23^. By integrating these enzymatic H_2_S generation strategies with the genetically encoded biosensor hsGFP^13^, we establish a system that enables both dynamic production and real-time visualization of intracellular H_2_S (**Figure 1**).

**Figure 1.**
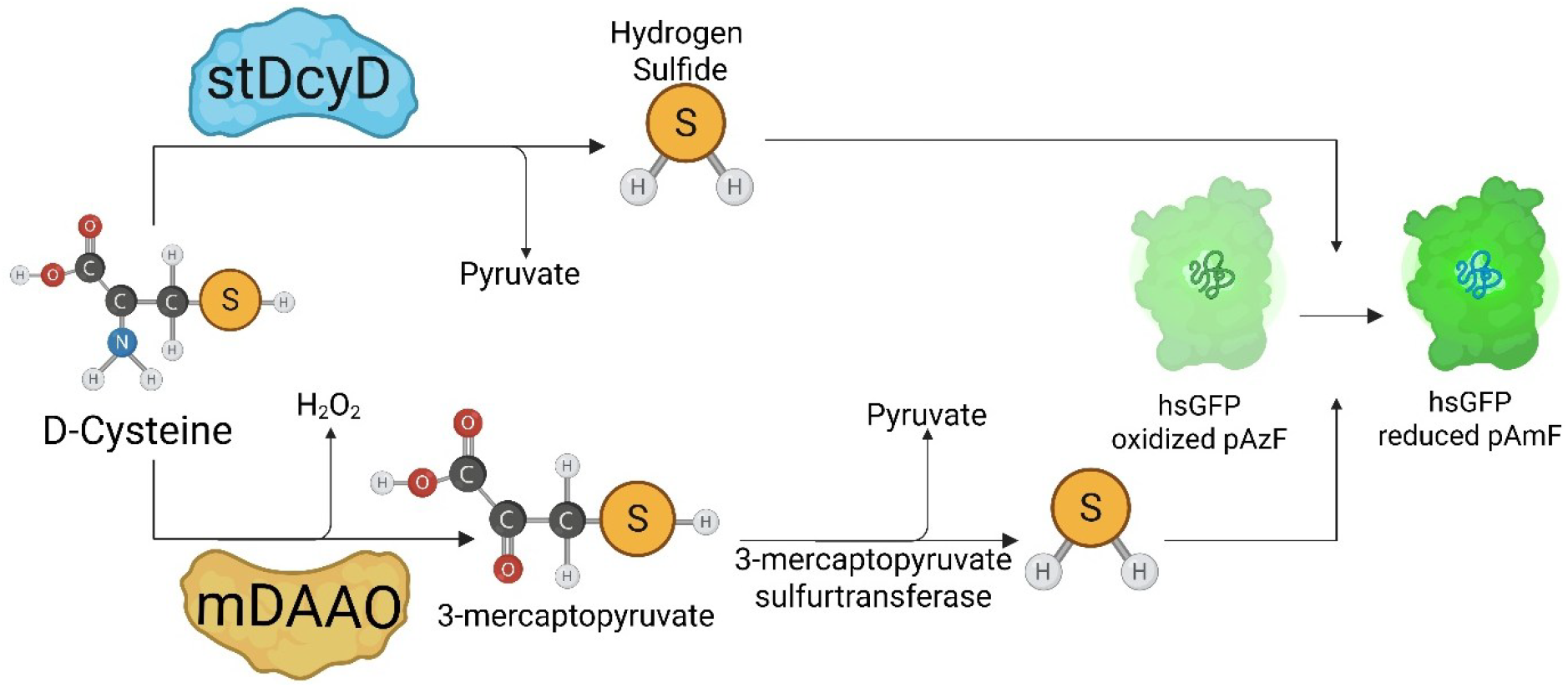
Schematic representation of two chemogenetic strategies for H_2_S generation. The bacterial enzyme stDCyD directly metabolizes D-cysteine into H_2_S, which can be detected using the genetically encoded H_2_S sensor, hsGFP. Alternatively, the modified D-amino acid oxidase (mDAAO) first converts D-cysteine into 3-mercaptopyruvate (3-MP) while generating H_2_O_2_ as a byproduct. The endogenous enzyme mercaptopyruvate sulfur transferase (3-MPST) subsequently converts 3-MP into H_2_S, which is also detectable by the hsGFP biosensor.

Given that both the endogenous enzyme MPST and stDCyD generate pyruvate as a byproduct during H_2_S production^**17**,**24**^, we sought to assess pyruvate formation as an indirect measure of enzymatic activity and efficiency. To this end, we employed the genetically encoded pyruvate biosensor Pyronic^**25**^ for real-time monitoring of intracellular pyruvate levels (**Figure 2A**). Upon administration of 5 mM D-cysteine, real-time imaging revealed no significant changes in pyruvate levels in either wild-type (WT) cells or cells expressing mCherry-mDAAO (**Figure 2B,C**). In contrast, cells expressing DsRed-stDCyD exhibited an increase in pyruvate generation (**Figure 2C, middle panel**), indicating a higher catalytic efficiency of stDCyD in metabolizing D-cysteine. Since the production of H_2_S and pyruvate by stDCyD occurs in equimolar amounts, these findings further support the suitability of stDCyD as an effective tool for controlled H_2_S generation.

**Figure 2.**
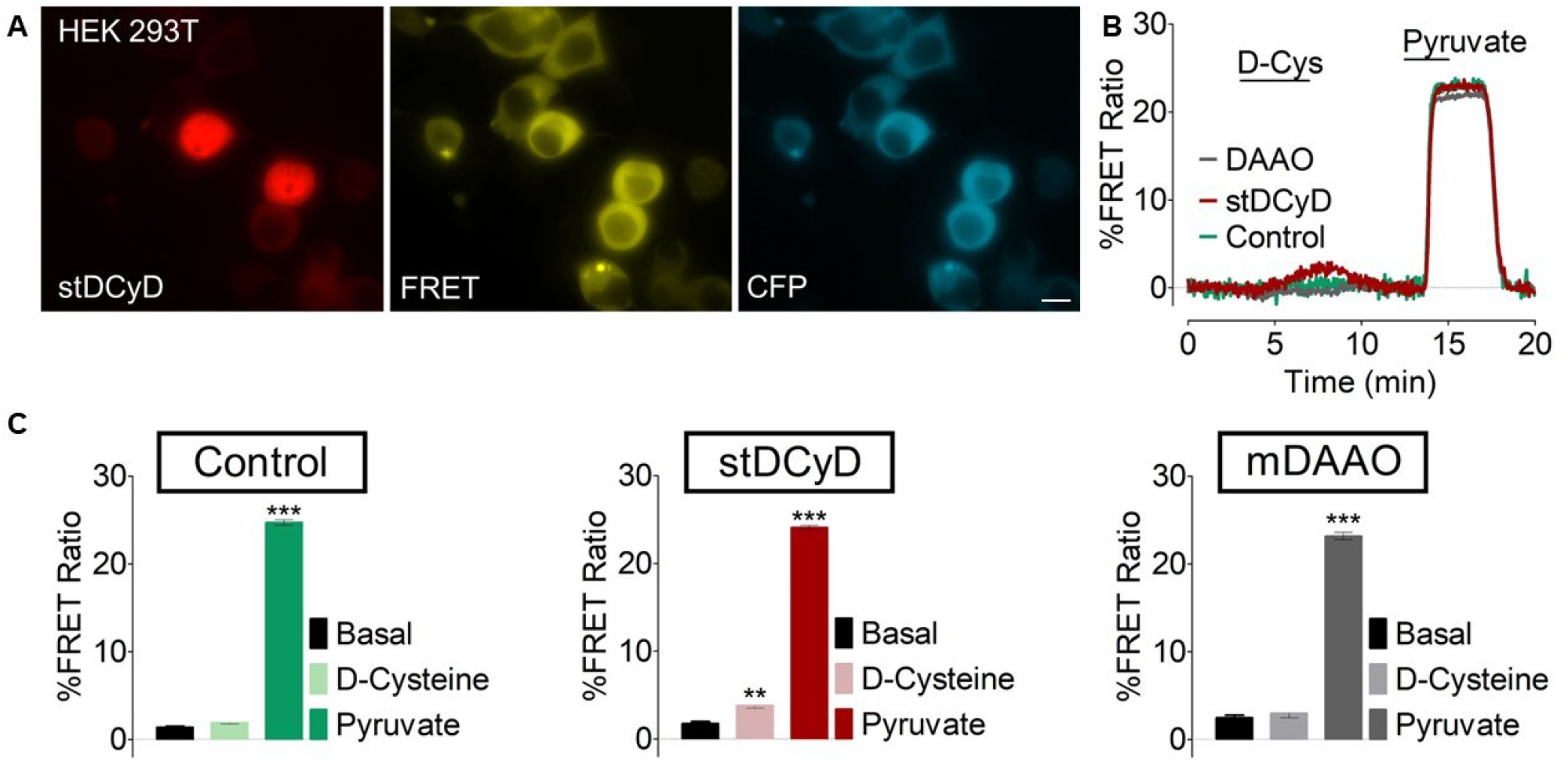
Pyruvate as a byproduct of enzymatic activity of stDCyD. **A)** Wide-field images of HEK 293T cells co-expressing DsRed-stDCyD (left panel), and Pyronic biosensor (middle and the right panels), scale bar represents 50 μm. **B)** Real-time traces of Pyruvate generation in HEK 293T cells co-expressing DsRed-stDCyD and Pyronic (red curve, 3/29) versus cells co-expressing mCherry-mDAAO and Pyronic (grey curve, 3/34), and cells only expressing Pyronic (green curve, 3/41) in presence of 5 mM D-Cysteine. At the end of the experiment cells were perfused with 5 mM pyruvate. **C)** One-way ANOVA analysis of FRET ratio in response to 5 mM D-cysteine and 5 mM pyruvate relative to the basal ratio.

Previous work, including our own, has established that mDAAO generates hydrogen peroxide (H_2_O_2_) during the oxidation of D-amino acids to the2ir corresponding α-keto acids^**26**,**27**^. To directly compare H_2_O_2_ production between mDAAO and stDCyD, we co-transfected HEK 293 cells with plasmids encoding either mCherry-mDAAO or DsRed-stDCyD, together with the ratiometric H_2_O_2_ biosensor HyPer7.2^**28**^. Upon acute administration of 5 mM D-cysteine, real-time imaging revealed a robust increase in intracellular H_2_O_2_ levels in cells expressing mCherry-mDAAO, consistent with the enzyme’s oxidative activity. In contrast, cells expressing DsRed-stDCyD exhibited no detectable change in H_2_O_2_ levels under the same conditions. These findings demonstrate that, unlike mDAAO, stDCyD enables selective H_2_S generation without concomitant oxidative stress, positioning it as a superior chemogenetic tool for controlled intracellular H_2_S modulation. Based on these findings, we selected stDCyD for subsequent experiments.

As the next step, we aimed to directly visualize H_2_S generation resulting from stDCyD activity using the genetically encoded H_2_S biosensor, hsGFP^**13**^. WT HEK 293T cells and cells expressing stDCyD were transiently transfected with the hsGFP sensor. Upon acute administration of 10 mM D-cysteine, an increase in H_2_S levels was observed in both WT and stDCyD-expressing cells. However, the maximum response was significantly greater in cells expressing stDCyD compared to WT cells. These results provide direct evidence that stDCyD efficiently metabolizes D-cysteine to generate H_2_S and validate its utility as a robust tool for intracellular H_2_S production.

## Discussion

Similar to H_2_O_2_, H_2_S was historically regarded as a toxic molecule^**29**^. Only in recent years critical role of H_2_S in cellular physiology been recognized^**2**^. This shift in understanding has catalyzed significant efforts to develop both chemical and genetically encoded probes for the real-time visualization of this gasotransmitter *in vitro* and *in vivo*. Current literature indicates that H_2_S exerts both beneficial and harmful effects depending on its concentration and the duration of exposure. At low concentrations, H_2_S supports various cellular processes, including signaling, energy metabolism, and protection against oxidative stress. However, at higher concentrations, it can exert cytotoxic effects and contribute to oxidative damage^**3**^. Thus, the ability to dynamically and precisely manipulate intracellular H_2_S levels represents an important unmet need in the field.

Existing approaches to modulate H_2_S levels include the genetic manipulation of endogenous enzymes—such as CBS, CSE, and MPST—the application of H_2_S-releasing chemical cages, and the administration of sulfide salts (e.g., sodium hydrosulfide, NaHS). However, these strategies have notable limitations, including irreversibility, poor suitability for *in vivo*^**30**^ use, off-target effects^**20**^, and a lack of spatiotemporal precision. Therefore, in this study, we have established a more efficient, reversible, and non-invasive method for the controlled generation of intracellular H_2_S.

We evaluated two candidate enzymes for this purpose: mDAAO and stDCyD. In our previous work, we demonstrated that mDAAO can catalyze H_2_S production in the presence of D-cysteine^**16**^. Independently, the literature has also reported that stDCyD can generate H_2_S using D-cysteine as a substrate^**17**^. However, the mechanisms by which these enzymes produce H_2_S are distinct (**Figure 1**). stDCyD directly converts D-cysteine into H_2_S and pyruvate, utilizing PLP as a cofactor. In contrast, mDAAO first oxidizes D-cysteine into its corresponding α-keto acid, mercaptopyruvate, which is subsequently metabolized to H_2_S by endogenous MPST.

Both MPST and stDCyD produce pyruvate as a byproduct. Therefore, to indirectly compare the efficiency of these two enzymatic pathways (as well as the endogenous pathway), we used the genetically encoded pyruvate biosensor Pyronic to monitor pyruvate production in real time (**Figure 2**). Our results showed that significant pyruvate generation occurred only in cells expressing stDCyD, but not in WT cells or those expressing mDAAO. These findings suggest that stDCyD exhibits superior catalytic efficiency for D-cysteine metabolism under the tested conditions.

Further characterization of the enzymatic activities focused on H_2_O_2_ generation. It is well established that mDAAO, while catalyzing the oxidation of D-amino acids, produces H_2_O_2_ as a byproduct. Using the ratiometric H_2_O_2_ sensor HyPer7.2, we observed a robust increase in intracellular H_2_O_2_ following acute administration of D-cysteine in cells expressing mDAAO. In contrast, cells expressing stDCyD showed no detectable H_2_O_2_ production under the same conditions (**Figure 3**). This distinction further highlighted the advantage of stDCyD for specific H_2_S manipulation without the confounding effects of oxidative stress induced by H_2_O_2_. Based on these observations, we selected stDCyD as the preferred chemogenetic tool for subsequent experiments.

**Figure 3.**
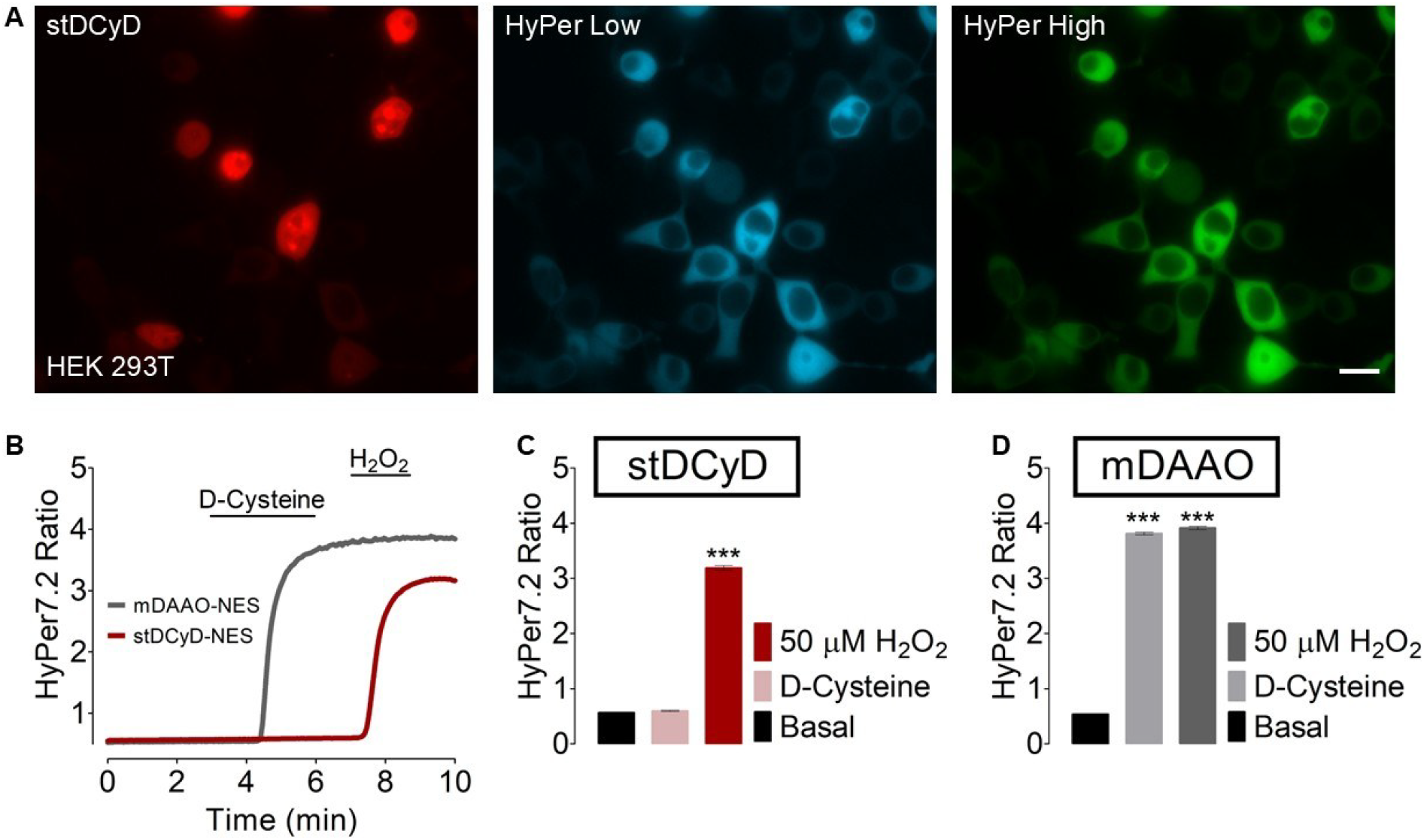
mDAAO, but not stDCyD, generates H_2_O_2_ in the presence of D-cysteine. **A)** Wide-field images of HEK 293T cells co-expressing DsRed-stDCyD (left panel) and the HyPer7.2 biosensor (middle and right panels). Scale bar: 50 μm. **B)** Real-time traces of H_2_O_2_ generation in HEK 293T cells co-expressing DsRed-stDCyD and HyPer7.2 (red curve, 3/36) versus cells co-expressing mCherry-mDAAO and HyPer7.2 (gray curve, 3/40) upon addition of 5 mM D-cysteine. At the end of the experiment, cells were perfused with 50 μM exogenous H_2_O_2_. **C, D)** One-way ANOVA analysis of HyPer7.2 ratio changes in response to 5 mM D-cysteine and 50 μM H_2_O_2_ relative to the basal ratio.

To directly visualize H_2_S generation by stDCyD, we employed the genetically encoded H_2_S biosensor, hsGFP. Acute administration of 10 mM D-cysteine resulted in a significant increase in the hsGFP signal in cells expressing stDCyD, confirming active H_2_S production. Interestingly, a modest H_2_S signal was also detected in WT cells upon D-cysteine treatment (**Figure 4**). Two plausible explanations for this observation exist based on prior reports. First, endogenous D-amino acid oxidase may oxidize D-cysteine to mercaptopyruvate, which is then converted to H_2_S by MPST^**23**^. Second, D-cysteine may be racemized to L-cysteine by endogenous serine racemase^**31**^, after which it can be metabolized by canonical H_2_S-producing enzymes such as CBS and CSE. To determine the contribution of these pathways, we plan to generate knockout (KO) cell lines targeting key enzymes involved in these alternative routes.

**Figure 4.**
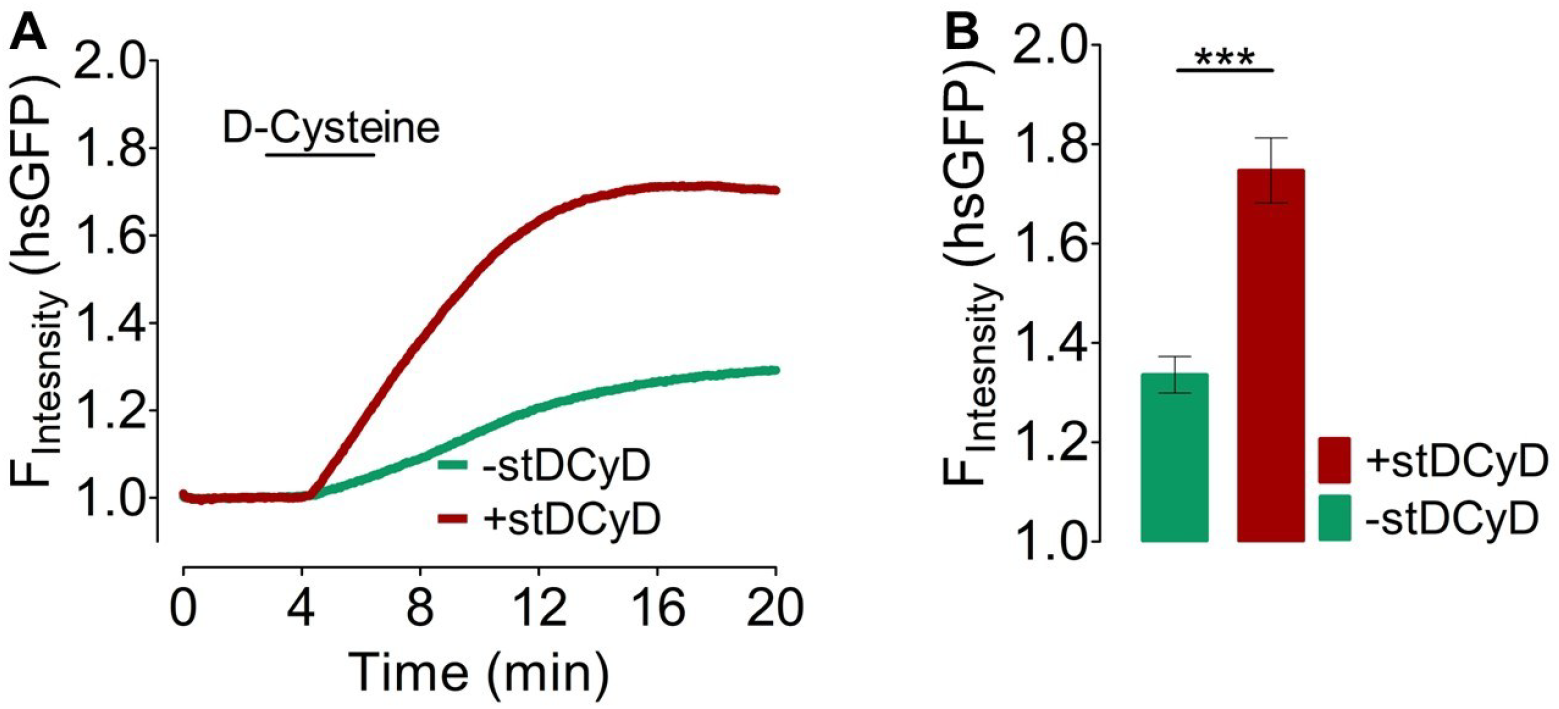
H_2_S generation in HEK 293T cells upon enzymatic activity of stDCyD. **A)** Panel shows representative real-time traces of produced H_2_S in response to 10 mM D-Cysteine in HEK 293T cells co-expressing untargeted hsGFP and stDcyD (red curve, 3/41) or only hsGFP (green curve, 3/54). **B)** The bar graph shows statistical analysis of the maximum response of hsGFP in response to 10 mM D-Cysteine.

## Conclusion

Our study introduces stDCyD as a robust, reversible, and precise tool for the intracellular generation of H_2_S, overcoming several limitations associated with existing methods. This platform will enable new investigations into the complex and dynamic roles of H_2_S in cellular physiology and pathophysiology.

## Materials and Methods

### Cell Culture

Characterization experiments were conducted using cultured human embryonic kidney (HEK 293 T) cells maintained in a high-glucose (4.5 g/L) complete medium supplemented with 10% fetal bovine serum (FBS), 100 μg/ml streptomycin, and 100 U/ml penicillin, incubated at 37°C with 5% CO_2_. Approximately 24 hours before transfection, cells were plated onto 30 mm No.1 glass coverslips (Glaswarenfabrik Karl Knecht Sondheim, Germany). Once the cells reached about 70– 80% confluency, they were co-transfected with the appropriate mammalian expression plasmids using PolyJet transfection reagent, following the manufacturer’s protocol. Imaging was carried out 24 hours post-transfection.

### H_2_S imaging using hsGFP biosensor

For H_2_S detection, cells were seeded onto 30 mm glass coverslips approximately 24 hours prior to transfection. Upon reaching ∼70–80% confluency, cells were co-transfected with the pMAH-POLY and the hsGFP biosensor plasmids using PolyJet transfection reagent. Five hours post-transfection, the culture medium was replaced with L-methionine and L-cystine-free DMEM supplemented with 2 mM *p*-azidophenylalanine (*P*AZF) to enable non-canonical amino acid incorporation. After 24 hours of incubation, residual *P*AZF was removed by exchanging the medium for standard DMEM lacking L-methionine and L-cystine. Imaging experiments were conducted 24 hours following the *P*AZF washout.

### Buffers and Solutions

Unless otherwise stated all chemicals were purchased from NeoFroxx. To maintain cells outside of the cell culture incubator, a cell storage buffer containing 2 mM CaCl_2_, 5 mM KCl, 138 mM NaCl, 1 mM MgCl_2_, 10 mM HEPES, 0.44 mM KH_2_PO_4_, 2.6 mM NaHCO_3_, 0.34 mM NaH2PO4, 10 mM D-Glucose, 0.1% MEM Vitamins (Pan-Biotech, Aidenbach, Germany), 0.2% essential amino acids (Pan-Biotech, Aidenbach, Germany), 100 μg/mL Penicillin (Pan-Biotech, Aidenbach, Germany), and 100 U/mL Streptomycin (Pan-Biotech, Aidenbach, Germany) was used. The pH was adjusted to 7.42 using 1 M NaOH. For live-cell imaging experiments, a HEPES buffer solution was used which consisted 2 mM CaCl_2_, 5 mM KCl, 138 mM NaCl, 1 mM MgCl_2_, 10 mM HEPES, 10 mM D-Glucose, and pH was adjusted to 7.42 using 1 M NaOH.

### Widefield epifluorescence microscopy

Widefield epifluorescence microscopy was conducted using the Axio Observer Z1.7 inverted system (Carl Zeiss AG, Oberkochen, Germany), featuring a high-resolution Plan-Apochromat 40×/1.4 numerical aperture oil immersion lens and a monochrome Axiocam 503 CCD camera. A custom-built, gravity-based perfusion apparatus was integrated to enable precise delivery and removal of experimental solutions. For visualization of HyPer7-expressing cells, excitation was alternated between 423/44 nm and 469/38 nm wavelengths via a motorized filter wheel fitted with FT455 and FT495 dichroic beamsplitters (HyPer Low and High channels, respectively). Emission signals were collected through a common bandpass filter (525/50 nm) to facilitate ratiometric imaging. Cells expressing mcherry-mDAAO or DsRed-stDyCD were illuminated with 555/30 nm light, and emitted fluorescence was filtered through an FT570 dichroic mirror and 605/70 nm bandpass filter. FRET-based pyruvate sensor imaging was performed using 430 nm excitation, with emissions monitored at 480 nm and 525 nm for CFP and FRET channels respectively. Detection of H_2_S levels with the hsGFP biosensor was achieved by exciting at 480 nm and capturing emission at 525 nm. Image acquisition and system control were executed using Zen Blue 3.1 Pro software (Carl Zeiss AG).

### Statistical Analysis

Imaging data were processed using GraphPad Prism version 5 (GraphPad Software, San Diego, CA, USA). Each experiment was repeated a minimum of three times, with results reported as N/n—where N denotes the number of independent experiments and n the number of cells analyzed. Data are presented as mean ± SEM, unless specified otherwise. For comparisons involving more than two groups, statistical significance was determined using one-way ANOVA followed by Dunnett’s post hoc test (comparing each group to the control). For comparisons between two groups, an unpaired Student’s t-test was applied.

## Acknowledgments

Figure 1 was generated with BioRender.com (Agreement Number: BT289BFFTZ)

